# Pandemic, epidemic, endemic: B cell repertoire analysis reveals unique anti-viral responses to SARS-CoV-2, Ebola and Respiratory Syncytial Virus

**DOI:** 10.1101/2021.08.19.456951

**Authors:** Alexander Stewart, Emma Sinclair, Joseph Ng, Joselli Silvia O’Hare, Audrey Page, Ilaria Serangeli, Christian Margreitter, Nora Kasar, Katherine Longman, Cecile Frampas, Catia Costa, Holly Lewis, Bryan Wu, David Kipling, Peter Openshaw, Christopher Chu, J Kenneth Baillie, Janet T Scott, Malcolm G Semple, Melanie Bailey, Franca Fraternali, Deborah Dunn-Walters

## Abstract

Immunoglobulin gene heterogeneity reflects the diversity and focus of the humoral immune response towards different infections, enabling inference of B cell development processes. Detailed compositional and lineage analysis of long read IGH repertoire sequencing, combining examples of pandemic, epidemic and endemic viral infections with control and vaccination samples, demonstrates general responses including increased use of *IGHV4-39* in both EBOV and COVID-19 infection cohorts. We also show unique characteristics absent in RSV infection or yellow fever vaccine samples: EBOV survivors show unprecedented high levels of class switching events while COVID-19 repertoires from acute disease appear underdeveloped. Despite the high levels of clonal expansion in COVID-19 IgG1 repertoires there is a striking lack of evidence of germinal centre mutation and selection. Given the differences in COVID-19 morbidity and mortality with age, it is also pertinent that we find significant differences in repertoire characteristics between young and old patients. Our data supports the hypothesis that a primary viral challenge can result in a strong but immature humoral response where failures in selection of the repertoire risks off-target effects.

## Introduction

The emergence of SARS-CoV-2 in 2019, the ensuing pandemic and evolution of novel variants continues to make COVID-19 a matter of global public health significance. The recent SARS, MERS, Zika and Ebola outbreaks have also highlighted a need to better understand how the human immune system responds to novel infections, develop better treatments and control their emergence and spread. Initial reports from the COVID19 pandemic, relying heavily on serum antibody titres, saw rapid declines in SARS-CoV-2 specific antibodies^1^ that raised concerns over the nature and duration of B cell memory. While total antibody titres decrease the persistent presence of SARS-CoV-2-specific memory responses some months after infection mitigates these concerns^2,3^.

Immunoglobulins (Ig), both as secreted antibodies and as B Cell Receptors (BCRs), mediate immunity against multiple pathogens through their vast variability in antigen binding. This variability is produced by V-D-J recombination ^4^, where V, D and J genes are recombined from a pool of diverse genes. B cells with Ig genes encoding disease-specific antibodies are expanded upon challenge, causing a skewing of the repertoire towards greater use of antigen-specific genes associated with the challenge in question. Furthermore, the imprecise joining of gene segments, together with the action of terminal deoxynucleotidyl transferase (TdT) creates a highly diverse complementarity determining region (CDR)3 region, which is important for antigen binding, and can be used to identify “clones” of B cells within a repertoire. These clonal assignments allow us to track lineages and follow the progress of the post-activation diversification events of somatic hypermutation (SHM) and class switching (CSR) as the B cell response develops. Thus, repertoire analyses can help to characterise changes in the memory/effector B cell compartments and identify individual genes of interest for possible antibody therapeutics.

Both SHM and CSR are mediated by the enzyme Activation Induced cytidine Deaminase (AID) and have traditionally been associated with germinal centre events in secondary lymphoid tissue, involving T cell help ^5–7^. There is, however, also evidence that CSR may occur outside the germinal centre environment ^8–11^ and may not require direct T cell help. The ability of a B cell to mount a directed effector response prior to the formation of a germinal centre allows a more rapid immune response but with lower affinity.

Immune responses are often impaired in older people, which has been of particular concern in COVID-19 patients. The older immune system has shown reduced responses to vaccination, frequently with higher numbers of autoreactive antibodies and inflammatory cytokines^12–14^. In B cells we, and others, have shown that particular subsets of B cells are altered with age: IgM memory cells (CD19+CD27+IgD+) are decreased in older people while the Double Negative (CD19+CD27-IgD-) are increased ^15,16^. Since IgM memory cells are often associated with a T-independent response, the decrease in IgM memory in older people could have severe consequences in infections where a rapid extrafollicular response is required^17,18^. It has also been shown that the B cell repertoire is skewed towards sequences with longer more hydrophobic CDR3 regions as we age^16,19^. As an immune response can result in a shift towards lower, less hydrophobic CDR3 regions^14,20^, and higher hydrophobicity has previously been correlated with increased polyspecificity^21–23^, the older immune repertoire seems to be disadvantaged in this respect.

In this study we took a long-read repertoire amplification approach that allowed us to track the V-D-J clonal lineages in the context of antibody subclass to better understand, compare and contrast B cell responses to emerging or endemic viruses. Samples were taken from COVID19 patients during and after infection, Ebola virus disease (EBOV) survivors from West Africa and the UK, volunteers challenged with Respiratory Syncytial Virus and compared with samples from healthy donors. We report the variation of repertoire between disease states in novel virus infection, with a focus on elderly who are known to respond less well to infection, particularly in SARS-CoV-2.

## Methods

### Sample collection

Whole blood samples (RSV, COVID19, Healthy) were collected into Tempus^™^ Blood RNA tubes, kept at 4°C, and frozen down to -20°C within 12 hours. Ebola samples were cone filters from plasmapheresis, dissolved in Tri reagent. RNA was extracted using Tempus^™^ kits according to instructions. Healthy samples taken after SARS-CoV-2 emergence were all confirmed negative for anti-SARS-CoV-2 antibodies by SureScreen lateral flow test and by ELISA ^24^. Ebola RNA blood samples were collected from convalescent patients with viral RNA negative PCR tests in the 2014-2016 West African outbreak, three patients were Caucasian treated in the UK, and the remaining were convalescent plasma donor participants from a trial in Sierra Leone^25^ (consented under the Sierra Leone Ethics and Scientific Review Committee ISRCTN13990511 and PACTR201602001355272 and authorised by Pharmacy Board of Sierra Leone, #PBSL/CTAN/MOHS-CST001). COVID-19 samples were collected from SARS Cov 2 positive patients at Frimley and Wexham Park hospitals during 2020 (consented under UK London REC 14/LO/1221). Each participant was attributed a “severity score” in relation to their fitness observations at the time of hospital admission using the metadata collected. This score used the “mortality scoring” approach of SR Knight *et al*. adapted to disregard age, sex at birth and comorbidities, and ranged from 0 to 6; patients scoring 0 to 3 were attributed low severity and patients scoring 4 to 6 were attributed high severity^25^. Convalescent COVID-19 patients, from hospital sampling, were contacted for further donations and sample taken 2-3 months post hospital discharge. RSV samples were collected from participants who took part in a human challenge study and were monitored for infection by viral PCR tests (consented under UK London REC 11/LO/1826). Briefly, healthy participants were challenged intranasally with 10^4^ plaque-forming units of the M37 strain of RSV and monitored for up to 6 months as previously described ^26^.

### Repertoire Library Generation

Tempus tube samples were defrosted at room temperature and RNA was extracted using the Tempus RNA extraction kit according to the manufacturer’s instructions. RNA samples were template switch reverse transcribed using SMARTScribe^™^ reverse transcriptase (Clonetech) according to manufacturer’s instructions using the SmartNNN TSO Primer (Supp. Methods Table 1) with a minimum of 170 ng of RNA input. The sample was then treated with 0.5 units/μl of Uracil-DNA Glycosylase (NEB) for 60 min at 37°C to reduce UMI interference, then incubated at 95°C for 10 min to inactivate the enzyme. Samples were amplified using Q5 polymerase (NEB) according to manufacturer’s instructions with an annealing step of 65°C for 20s and extension step of 72°C for 50 s for 21 cycles. Round one of PCR was performed with forward primer Smart20 and mixed heavy chain (IG[M, G, A]-R1) reverse primers (Supp. Methods Table 1). For PCR1 8 ⨯ 20 μl reactions were performed with 1 μl of RNA input per reaction. A semi-nested 2nd PCR was performed with forward primer PID-Step and reverse primers IG[M/G/A]-R2 (Supp. Methods Table 1); 16 reactions of 20μl each was performed for each isotype with the same thermal cycling conditions as PCR1 but with 12 cycles, with 1μl of template. The primers in PCR2 also contain Patient Identifier (PID) sequences to allow multiplexing on PacBio (Supp Methods Table 2). Samples were run on a bioanalyzer (Agilent 7500), isotypes from patients were pooled at equal concentrations and concentrated using Wizard PCR Clean-up kits (Promega) according to manufacturer’s instructions with 30 μl of elute. Each isotype was then purified using a PippinPrep^™^ with Marker K reagents (Sage Biosciences) used as an external ladder reference (IgM/G/A 600-100bp). The concentration was checked using a DNA quantification kit on the Qubit according to manufacturer’s protocol, the different isotype samples were pooled at equal concentrations and purified with SPRIselect beads (Beckman Coulter) at X0.8 sample volume with elution in 30μl of TE buffer. Sequencing was performed on either the PacBio RSII or Sequel platforms (See Supp. Methods Table 2).

Quality control, data cleaning and removal of multiplicated UMIs was carried out as previously published^16,27^. Immunoglobulin V-D-J gene usage and CDRH3 was determined using IMGT/High V-quest. Clonotype clustering was carried out as per^16,27^, in brief: a Levenshtein distance matrix was generated on the CDRH3, hierarchically clustered and branches cut at 0.05 to generate clones. Physicochemical properties were calculated using the R Peptides package ^28^.

### Analysis of clonal diversity

We sought for methods to qualitatively (visualising clone size distribution) and quantitatively compare clonal diversity (calculating metrics which summarise clonal diversity). We first noted, as one would expect, that sequencing depth (i.e., number of sequences sampled per repertoire) was a strong predictor of the number of clones (Supplementary Figure S1). For all repertoires considered here, a wide range of sequencing depth was observed (number of sequences range from 836 to 105,323, median = 12,040). We therefore adopted the following procedure in this analysis: first, to quantify the extent of clonal diversity we used the Gini coefficient which measures the evenness in the distribution of clone size across clones; application of this metric to quantify BCR clonal diversity has been well documented ^29–31^. For a given repertoire, clones were ordered by their clone sizes, and the cumulative distributions of clone sizes (in terms of percentage of sequences in repertoire) and its percentile distribution were compared for evenness. As such, the resulting metric was independent from the *absolute* numbers of sequences and clones, thereby allowing fair comparison across repertoire of different sequencing depths. As Gini coefficient is an indicator of evenness, we took (1 – Gini coefficient) as the metric of clonal diversity. To qualitatively compare clonal diversity, we generated visualisation using the following procedure to minimise the impact of sequencing depth differences: we first sampled 12,000 sequences (≈ median sequencing depth; see above) from each repertoire; for repertoires with less than 12,000 unique observations, this number of sequences were sampled with replacement. We then sampled up to 100 clones with probability scaled by clone sizes to generate bubble plots where each bubble represents a clone and bubble sizes are scaled with clone sizes. Genotypic features like V gene usage can be represented as colours. Such plots were included in Figure 2c.

### Analysis of BCR clone lineage trees

Lineage trees were reconstructed using the maximum parsimony method implemented in the dnapars executable in the phylip package^32^. All clones with at least 3 sequences were considered; IMGT-gapped, V-gene nucleotide sequences of all observations in the clone together with the annotated germline V-gene sequence (included to root the tree) were included as input to dnapars. Functionalities implemented in the alakazam R package^33^ were used to call dnapars and reformat the output into text-based tree files (newick format) and directed graphs (igraph objects manipulated in R). The directed graphs were further parsed using functions in alakazam and igraph to obtain, for each observed sequence in the given clone, its distance *D* from the given germline gene *g* (denoted here as *D*_*g*_), as estimated by the dnapars-reconstructed lineage tree: the closer this distance is to 0, the closer the sequence is to germline and therefore a lower mutational level.

We sought to summarise, for a given clone, the distribution of *D*_*g*_; this distribution would indicate the overall mutational level of sequences within the clone (summarised using conventional statistics like the median of *D*_*g*_) and the evenness of mutational level (i.e. whether the clone consists of expansion of sequences with a similar mutational level, or it comprises sequences with a wide range of mutational levels). This can be visualised as a heatmap (clones [vertical axis] versus *D*_*g*_ [horizontal axis], with colours scaling with density of the distribution; see Figure 5b), or as a curve (clones [vertical axis] versus the median of *D*_*g*_ [horizontal axis], See Figure 5c). The curve representation allows calculation of area-under-curve (AUC) as a metric which we termed “Germline Likeness”, to quantify mutational levels across clones. This is similar to quantifying sequence similarity to germline, except that here Germline Likeness quantifies the tendency to which all clones from the BCR repertoire of a given individual have high similarity to the germline (and therefore lower mutational levels).

### Detecting class-switch recombination events from lineage trees

Since the lineage trees were constructed using only V-gene sequences (see above), in theory antibody sequences of different subclasses could be ordered in the tree in a way that imply class-switch recombination (CSR) events which are mechanistically impossible. We therefore pruned the dnapars-reconstructed tree to remove edges which imply CSR events that violates the physical order of constant region genes in the human IGH locus. This was performed using a Python implementation of the Edmond’s algorithm to construct a minimum spanning arborescence tree with the given germline V gene sequence as root. With this arborescence tree the type of CSR (subclass switched from/to) and the distance-from-germline at which the CSR event occurred (estimated as the median distance-from-germline of the two observations relevant to the given event) were obtained.

### Convergent network

Productive heavy chain sequences with CDRH3 of length shorter than 30 amino acids were considered in the construction of a convergent network. Sequences were connected if they meet the following criteria (a) same V and J gene usage; (b) from different individuals; (c) same CDRH3 amino acid length, and (d) **≥**85% CDRH3 amino acid identity. To allow interpretation of possible targets of sequences in convergent network clusters, known binders were included in constructing the network. Known binders were taken from the following sources: first experimentally determined antigen-antibody structural complexes deposited in the Protein Data Bank (PDB). PDBe was queried on 19 May 2021 with the search term ‘Organism: Severe acute respiratory syndrome coronavirus 2’. The resulting list of PDB entries were overlapped with entries in the SAbDab structural antibody databases^34^ to obtain list of PDB complexes of antibodies and SARS-CoV-2 proteins. A total of 215 heavy chains from 186 structures were considered. Second, known binders validated in experiments where antibody variable regions were cloned and assessed for SARS-CoV-2 protein binding were taken from published work ^35–40^. All known binder sequences were annotated for V/J gene usage using either IMGT/High-VQuest (if DNA sequences were provided) or IMGT/DomainGapAlign (if only amino acid sequences were provided). Information regarding specificity (i.e. SARS-CoV-2 protein targets) were obtained from either supplementary data files in the cited publications or by visual inspection (for PDB structures). Supplementary Table S5 contains all known binder sequences included in this analysis. To construct the network, known binders were connected to one another and to repertoire sequences using the identical criteria mentioned above. In total 809 unique CDRH3 sequences were considered in constructing the convergence network. The resulting network contains 7500 sequences (7370 from repertoire, 130 known binders). Analogous convergent networks were constructed using the Ebola and RSV repertoire data, separately considered with respective known binders and Healthy individuals’ repertoire; the majority of clusters were formed mainly of sequences from Healthy donors absent of known binders^41–44^, although we were able to identify two convergent clusters of RSV-infected individuals with similar CDRH3 to known binders of the RSV fusion glycoprotein (Supplementary Figure S6). To investigate whether clusters shared across disease conditions exist, convergent networks were also constructed considering CV, RSV and EBOV repertoire and binder sequences altogether (Supplementary Figure S7). Supplementary Table S6 contains all convergent networks constructed, presented as list of pairwise sequences.

### Statistical analysis and Data visualisation

V-D-J gene usage for each patient was turned into a proportion to normalise for different numbers of sequences and allow for comparison. Gene usage analysis was performed in GraphPad Prism 8.4.3 using a two-way ANOVA with a Dunnett’s post hoc test. All other statistical analyses were performed in the R statistical computing environment (version 4.0.2). Data visualisation was performed using the R ggplot2 package and the following specialised R packages: visNetwork (for visualising convergent CDRH3 network clusters) and ggseqlogo (for visualising CDRH3 sequence logos). PDB structures were visualised using PyMOL (version 2.3.0). Histograms of CDRH3 length and hydrophobicity, as measured by Kidera factor 4, were constructed on the Brepertoire website^45^.

## Results

### Patient cohorts

*IGH* sequences, of total V-D-J plus ∼150-200 bp of C regions, were obtained from pandemic, epidemic and endemic diseases and stages along with 24 healthy controls across multiple age ranges (Figure 1, Table 1). This included: 16 hospitalised COVID-19 patients (CV19), 5 of these patients had follow-up convalescent samples (CV19-Recovered, hereafter CV19R), 12 Ebola convalescent plasma donors (EBOV), 12 participants challenged with RSV; 6 of whom became infected (RSV-I) and 6 of whom did not (RSV-U). Healthy Samples (Healthy) were a grouping of YFVD0, RSVD0 and samples taken as controls during the COVID pandemic (n=24). Numbers of sequences varied from 836 to 105,323, median = 12,040 per sample, *IGH* gene usage for each patient was expressed as a proportion of the total in order to normalise for differences in sequence numbers between different samples.

**Figure 1.**
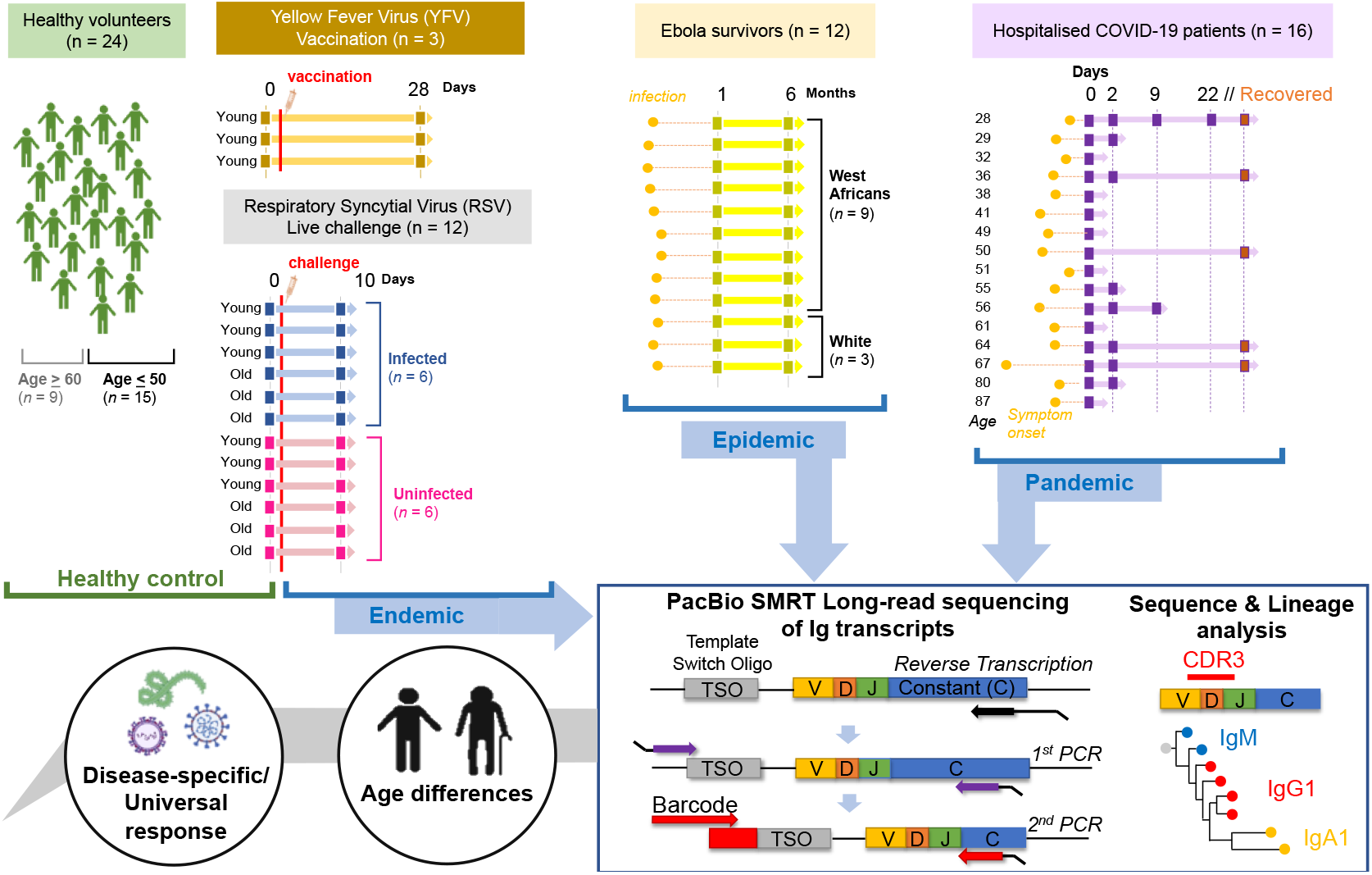
Schematic to illustrate data collection and analysis conducted in this study. Samples were taken from Healthy individuals, recovered Ebola survivors, hospitalised COVID-19 patients, live RSV challenge participants that either became infected or did not and Yellow Fever vaccine recipients before vaccine and 28 days post-inoculation. Extracted sample RNA was subject to a heavy gene specific race 5’ and nested PCR amplification process retaining V-D-J and sub-class information.

**Table 1.** Donor characteristics. See Supplementary Table S1 for a detailed summary of metadata per donor.

| Sample | Age | Gender | Ethnicity | COVID-19<br>Severity Score<br>(out of 6) | Days since<br>symptom onset |
| --- | --- | --- | --- | --- | --- |
| **Healthy**<br>**(n = 24)** | Median 29.5 (Range 23 - 76)<br>≤50 years old: 15/24 (62.5%)<br>≥60 years old: 9/24 (37.5%) | Female: 7/24 (29.2%)<br>Male: 5/24 (20.8%)<br>Unknown: 12/24 (50%) | White: 12/24 (50%)<br>Unknown: 12/24 (50%) |  |  |
| **COVID-19 (n = 16)** | Median 50.5 (Range 28 - 87)<br>≤50: 8/16 (50%)<br>50-60: 3/16 (18.75%)<br>≥60: 5/16 (31.25%) | Female: 7/16 (43.75%)<br>Male: 9/16 (56.25%) | White: 13/16 (81.25%)<br>South East Asian: 1/16 (6.25%)<br>Indian Subcontinent: 2/16 (12.5%) | Median 3 (Range 1 - 5) | Median 8 (Range 1 – 35) |
| **COVID-19 Recovered**<br>**(n = 5)** | Median 50 (Range 28 - 87)<br>≤50: 3/5 (60%)<br>≥60: 2/5 (40%) | Female: 3/5 (60%)<br>Male: 2/5 (40%) | White: 4/5 (80%)<br>Indian Subcontinent: 1/5 (20%) |  |  |
| **RSV Infected**<br>**(n = 6)** | Young: 3/6 (50%)<br>Older: 3/6 (50%) |  |  |  |  |
| **RSV Uninfected**<br>**(n = 6)** | Young: 3/6 (50%)<br>Older: 3/6 (50%) |  |  |  |  |
| **Ebola**<br>**(n = 12)** | Young: 3/12 (50%)<br>Unknown: 9/12 (50%) | Female: 1/12 (8.3%)<br>Male: 2/12 (16.7%)<br>Unknown: 9/12 (75%) | White: 3/12 (25%)<br>West African: 9/12 (75%) |  |  |
| **YFV D28 (n = 3)** | Median 28 (Range 27 - 28)<br>Young: 3/3 (100%) | Female: 1/3 (33.3%)<br>Male: 2/3 (66.7%) | White: 3/3 (100%) |  |  |

### IGH gene repertoire changes in response to viral infection

Although the humoral immune response is varied, with different subclasses of antibody having different effector functions^46^, many methods of repertoire analysis have hitherto not distinguished between antibody subclasses. We have used PacBio methods to obtain full V-D-J sequence in the context of subclass usage to investigate class switching events during immune responses to infection. We also distinguish between mutated versus unmutated IgM sequences, as a proxy for identifying IgM memory responses. Comparisons of subclass distribution, in relation to healthy controls, revealed a significant increase in the proportion of IgA1 compared to IgA2 in CV19, and RSV-I and the proportion of IgG1 relative to IgG2 in CV19, EBOV and RSV-I (Figure 2a, 2b). The differences in CV19 IgG and IgA repertoire returned to ‘normal’ healthy levels by the time of convalescent sampling (CV19-Recovered) 2-3 months later.

Immune challenge is characterised by clonal expansion of B cells that express Ig which reacts with the challenging antigen. We identify members of clones in the repertoire by clustering the CDRH3 regions and looking at the largest clones in each sample we can see evidence of increased clonal expansions in CV19 patients (Figure 2c). In the full CV19 repertoire *IGHV4* family genes were expanded (Figure 2c), more specifically of *IGHV4-39* (Supplementary Figure S2) and some *IGHV3* family, this is particularly noticeable in IgG1 and IgA1. Analysis of clonal diversity of memory B cells using the Gini index, taking all possible clones into account, found CV19 patients had a less clonally diversified repertoire in all but the IgG2 and IgG4 partitions (Figure 2d, Supplementary Figure S3), suggesting pervasive expansions of specific BCR clones. Unusually, we also saw a decrease in diversity of unmutated IgM sequences, indicative of clonal expansion prior to SHM and CSR (Figure 2d). These values returned to normal in the CV19R samples. In comparison, *IGHV1* family was expanded in the RSV-I IgG1 partition (Figure 2c), particularly of *IGHV1-18* (Supplementary Figure S4). Active infection with RSV showed an increase in diversity of IgA2, and samples taken 28 days after yellow fever vaccination showed an increase in diversity of both IgA2 and the mutated IgM populations. Interestingly, EBOV memory B cell populations were more diverse than healthy controls in all the main switched subclasses (IgG1, IgG2, IgA1, IgA2) (Figure 2d, Supplementary Figure S3).

**Figure 2.**
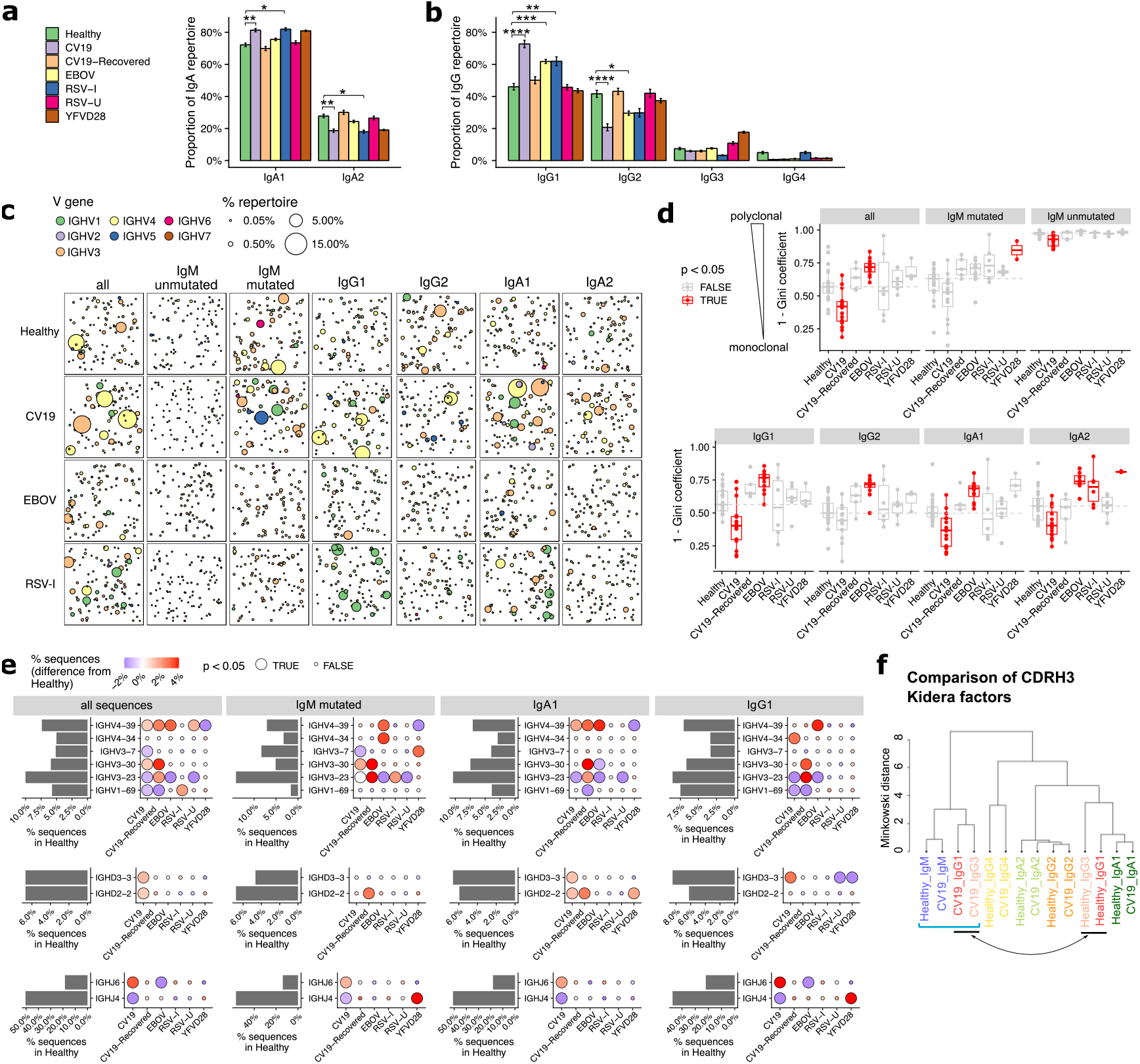
Distinct V-D-J and isotype usage in CV19, EBOV and RSV BCR repertoires. (**a-b**) Difference in sub-class use of IgA (A) and IgG (B) in viral disease and healthy BCR repertoires. (**c**) Clonal expansion of sequences of relevant effector types (as revealed in A) plus unmutated and mutated IgM to identify trends of V gene usage in viral infections. Each bubble sampled to a uniform depth (see Methods), with size proportional to clone size, represents one clone colour-coded by V-family usage. (**d**) Quantification of clonal expansion calculated using the Gini coefficient (see methods), revealed clonally expanded effector populations (more monoclonal/less diverse, closer to 1) or more diverse clones (closer to 0) in viral infections. Sample types with significant differences (p < 0.05) compared against Healthy were highlighted in red. Dashed line indicates the median diversity in the Healthy cohort. (**e**) Frequency of selected V-D-J gene usage in different cohorts for all sequences and further subdivided by IgM-mutated, IgA1, IgG1. Bar charts depict gene frequency usage in the Healthy cohort. Bubble plots depict the difference in usage (coloured: blue reduced/red increased) compared to healthy repertoires. (**f**) CDRH3 physicochemical characteristics (represented by Kidera factors) were analysed separated by sub-classes and disease status (Healthy/CV19), and compared using Minkowski distance. Note that IgG1 and IgG3 sequences from CV19 cluster together with IgM (square bracket), away from those of the same sub-classes from healthy individuals (indicated by arrow). Statistical significance in panels a, d and e was evaluated using one-way ANOVA and Dunnett post-hoc comparison against the Healthy cohort: p-value indicated either with colour (panel c), bubble size (e) or the symbols under the following scheme: *, p < 0.05; **, p < 0.01, ***, p < 0.001, ****, p < 0.0001.

Large clone sizes can mask whole repertoire changes, so we analysed the frequency of gene use after reducing the data to one representative sequence per clone. We found increased use of *IGHV3-30* in IgM mutated sequences in CV19 patients (Figure 2e), and also of IgM-mutated/IgA1/IgG1 in CV19R, this was unique to CV19. An increase in use of *IGHV4-39* was found in CV19 IgA1 sequences and was also found increased across the board in EBOV and RSV-U samples (Figure 2e). *IGHV3-23* was found to be reduced in ongoing infection (likely an offset as a result of relative increases in usage of other genes) but exceeded the healthy levels in CV19R and RSV-I. *IGHV1-69*, which has previously been associated with viral infections^47,48^ was increased in RSV-I but not EBOV or CV19. The YF day 28 vaccine samples increased use of *IGHV3-7* in IgM-mutated and *IGHV1-2* in IgG1 only (Supplementary Figure S4).

### Complementarity Determining Region 3 (CDRH3) immaturity in CV19

Given the importance of CDRH3 in antibody recognition, and the contribution to CDRH3 from the *IGHD* and *IGHJ* genes, we analysed these also. In CV19 samples there was a significant increase in use of *IGHD2-2, IGHD3-3* and *IGHJ6* (Figure2e). These genes tend to be more hydrophobic (*IGHJ6* being the exception) and all have among the longest amino acid lengths with only *IGHD3-16* being 2 AA longer. This contribution can be seen in the overall CV19 repertoire which skews towards longer amino acid sequences and increased hydrophobicity, indicative of early response as affinity maturation causes shorter less hydrophobic CDRs (Supplementary Figure S5). A clustering analysis of peptide physicochemical properties of CDRH3 regions generally results in a difference between IgM sequences and memory sequences (Figure 2f), presumably reflecting biases in antigen selection during post-challenge development. We can see that healthy and CV19 subclass sequences mostly have similar CDRH3 properties to each other, however, in the case of CV19 IgG1 and IgG3 cluster closer to IgM sequences from healthy and EBOV rather than healthy IgG1 and IgG3 sequences implying a more ‘naíve’, unselected, repertoire.

### Convergent antibody clusters between patients

To assess the functional importance of the skewed patterns of V, D and J gene usage in CV19 we created networks connecting sequences observed in our CV19 and control repertoire data (Figure 3a), using criteria previously employed in discovering ‘convergent’ antibody sequences shared between patients^49^. By also including known SARS-CoV-2 binders we obtain clusters of CDR3 sequences found in both CV19 patients and healthy controls, some of which converge towards known binders of SARS-CoV-2 proteins such as those targeting the receptor binding domain (RBD) of the spike protein(Figure 3b). Many of these large convergent clusters did not, however, include a known binder in the network (Figure 3c). Overall, convergent clusters use a diverse set of V genes, but most of our larger convergent clusters contain *IGHV3* or *IGHV4* families and demonstrate increased *IGHJ*6 usage as well as the more commonly used *IGHJ4* (Figure 3c). A comparison exclusively of the known binders to date reveals distinct combinations of V and J gene preferences (Figure 3d). We do find clusters of sequences using *IGHV3-53* and *IGHV1-58* such as those used in anti-RBD antibodies (e.g. Figure 3b). We find that sequences from convergent clusters tend to be found in larger clonal expansions than those without evidence of convergence (Figure 3e), possibly implying that specific clonal expansions in response to challenge are shared across patients. We note that half of the larger clusters have substantial contributions from healthy control sequences, so there may be some IGH genes, such as *IGHV3-33*/*IGHJ5* found also in RSV-I and EBOV convergent networks (Figure 3c, Supplementary Fig6a&b), which have increased versatility such that they are often seen in response to multiple different challenges.

**Figure 3.**
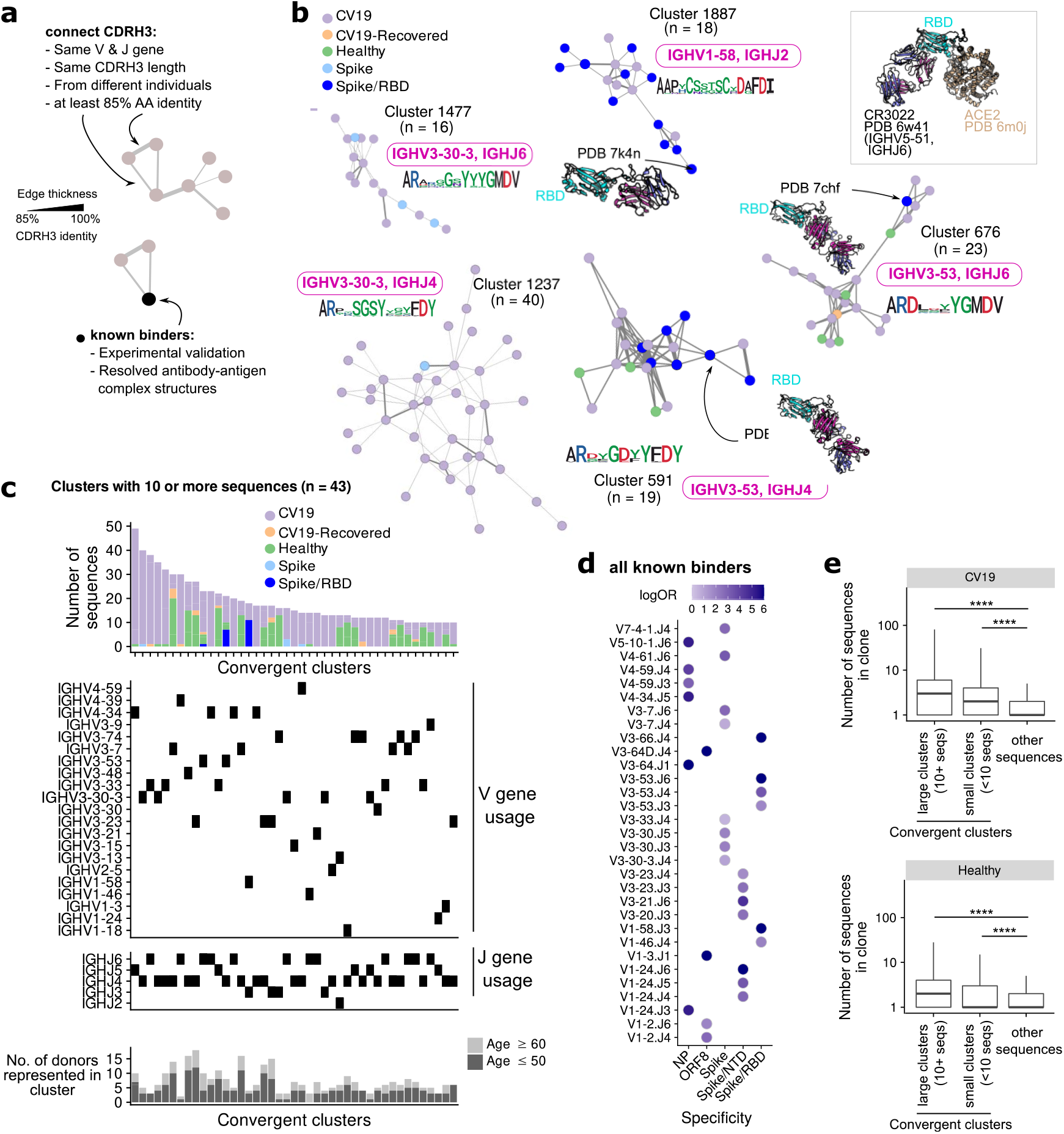
Convergent CDR3 clusters of sequences from CV19 repertoires and known SARS-CoV-2 binders. (**a**) CDR3 Known binder networks were created using same V, J and CDR3 length with at least 85% amino acid (AA) identity. (**b**) Convergent clusters from healthy and CV19 repertoire with known PDB structures. *IGHV* and *IGHJ* use and the CDR3 AA sequence were noted. (**c**) Clusters containing at least 10 sequences were visualized, with breakdown of repertoire origin (stacked bar plots), and the *IGHV* and *IGHJ* gene usage of each cluster aligned beneath. The number of donors with sequences in each depicted clusters are shown as bar graphs (bottom panel, **c**), broken down into subsets with age ≤50 (light grey) and ≥60 (dark grey). (**d**) All known binders were analysed for similarity of *IGHV*/J gene use to specific SAR-CoV-2 antibody targets (d). Dots coloured by enrichment (log-odds ratio, logOR) evaluated using Fisher’s exact test. Only V/J-specificity combinations with significant (p < 0.05) enrichment were shown. (**e**) Comparison of clonal expansion of convergent (split by clone size; ≥10 or <10 sequences) and non-convergent clusters in healthy and CV19 repertoires. Statistical significance evaluated using a Wilcoxon rank-sum test, ****: p < 0.0001. See Supplementary Figure S6 for analogous analyses on RSV and EBOV repertoires.

Similar analyses of RSV and EBOV repertoires were limited by the paucity of information on antibody binders, however it was notable that only RSV-I, and not RSV-U, showed evidence of convergence. *IGHV1-18* appears in a large cluster with a known RSV F-protein binder and although the large *IGHV3-23* cluster does not contain a known binder it forms part of the larger expansion of *IGHV3-23* genes in mutated IgM genes from this cohort (Figure 2e, Supplementary Figure S6b).

### Age-related differences

The disparity in CV19 severity and mortality between age groups is striking, so we looked for age-related differences in our B cell repertoire data. The difference in IgA1/IgA2 ratio is less in older people, not reaching significance. (Figure 4a). On the other hand, the increase usage of IgG1 in CV19 and concomitant decrease in IgG2 are robust across age (Figure 4b). Considering Ig gene usage, we observe the intriguing case of *IGHV3-30* which is only preferentially used by the over 60s during infection (Figure 4c). Conversely, *IGHV3-53*, which appears relatively frequently in known binder data in combination with *IGHJ4/6* but did not appear in our total cohort analysis (Figure 2e), is significantly increased in the under 50s IgM-mutated partition (Figure 4c). We also found that *IGHV4-39, IGHD2-2, IGHD3-3* and *IGHJ6*, which we find are expanded in CV19 across multiple B cell partitions, only have significantly increased expression in the under 50s and not the over 60s *IGHV4-34* appeared increased in both age groups. (Figure 4c).

**Figure 4.**
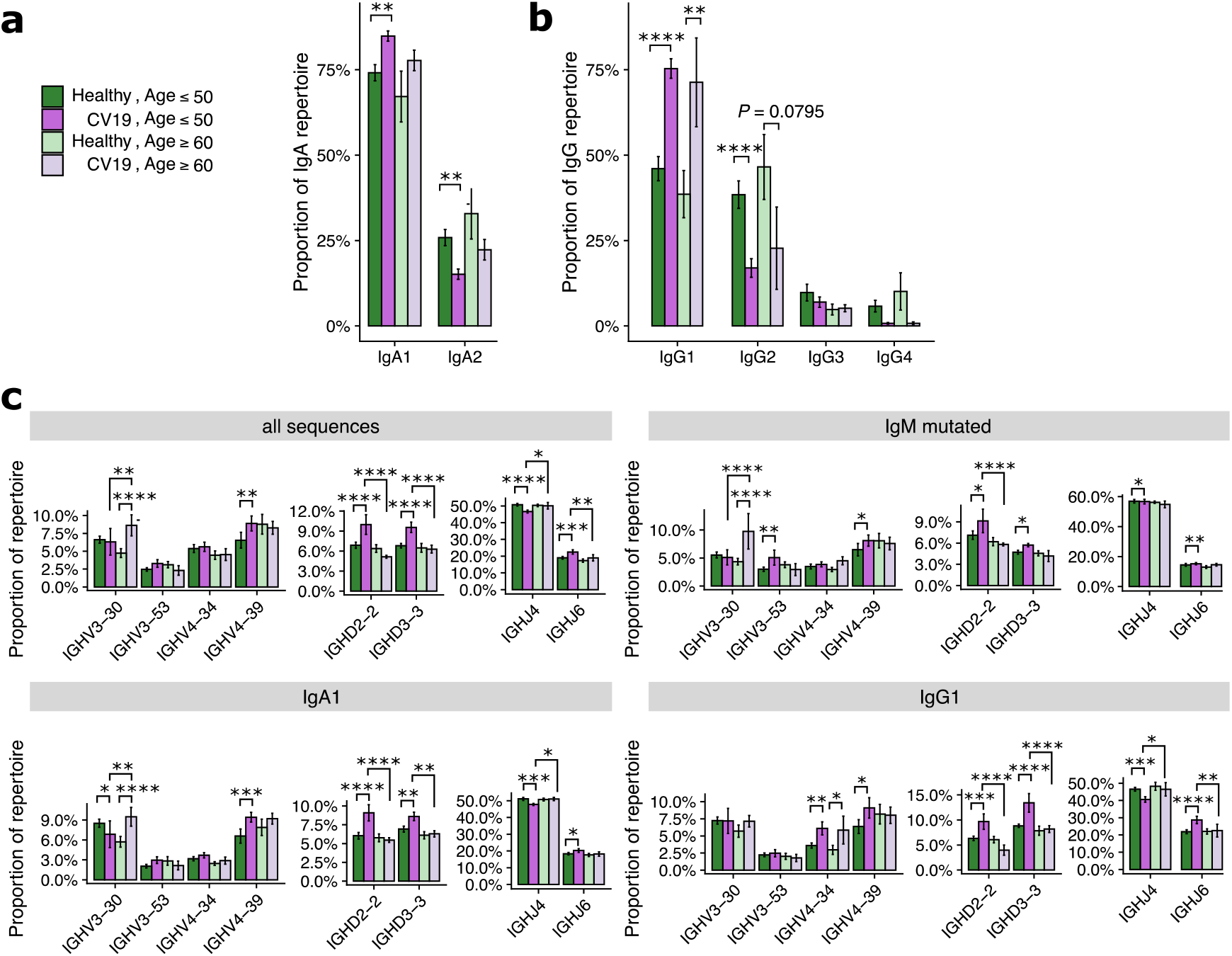
Age differences in V-D-J and isotype usage in CV19 repertoires. CV19 and healthy patients were split by over 60s and under 50s and were compared for IgA (**a**), IgG (**b**) usage and selected V-D-J gene usage (**c**). Statistical significance evaluated using two-way ANOVA and Tukey’s post-hoc test: *, p < 0.05; **, p < 0.01, ***, p < 0.001, ****, p < 0.0001..

### Immature IgG1 responses to CV19

Beyond the scope of gene usage, our BCR repertoire data also enabled reconstruction of individual BCR lineage trees to make inferences about the evolution of a particular clone. Using the annotated germline V allele as the root of the tree, we estimate, for each sequence in the lineage, its distance from the root (Figure 5a); this distance being directly proportional to mutational level. We visualise the distribution of this germline distance across all clonotypes observed in each given individual, and observe that the repertoire is dominated by clonotypes with very low mutational levels for a subset of CV19 patients, whilst the predominance of such clones is broadly absent in healthy controls (Figure 5b, c). Interestingly, in repertoires from convalescent individuals (both EBOV and CV19), we instead observe dominance of clonotypes with higher mutational levels, although the pattern is less striking than the CV19 patients during hospitalisation (Figure 5c). These curves allow for quantification of the Area Under the Curve (AUC), which constitutes a metric we term “Germline Likeness”: a higher Germline Likeness corresponds to a lower level of mutation across all clones (Figure 5d); this is akin to quantifying sequence similarity to the germline, except that Germline Likeness here quantifies such phenomenon for a given repertoire in general, rather than a specific sequence. Using this metric we confirm that CV19 repertoires were dominated by clones that were largely unmutated, while EBOV samples carried the greatest mutation rate (Figure 5e). As might be expected, with time to generate a germinal centre response, Germline Likeness in CV19 faded with time (Figure 5f), to the point where the CV19R repertoires have similar level of mutations compared to the EBOV-convalescent and healthy control repertoires. Partitioning the analysis by isotype, RSV and healthy controls demonstrate the expected trend where an increased level of mutations can be found in both IgG and IgA compared to IgM (Figure 5g). However, in CV19 only IgA showed a significant change in Germline Likeness from IgM (Figure 5g).

**Figure 5.**
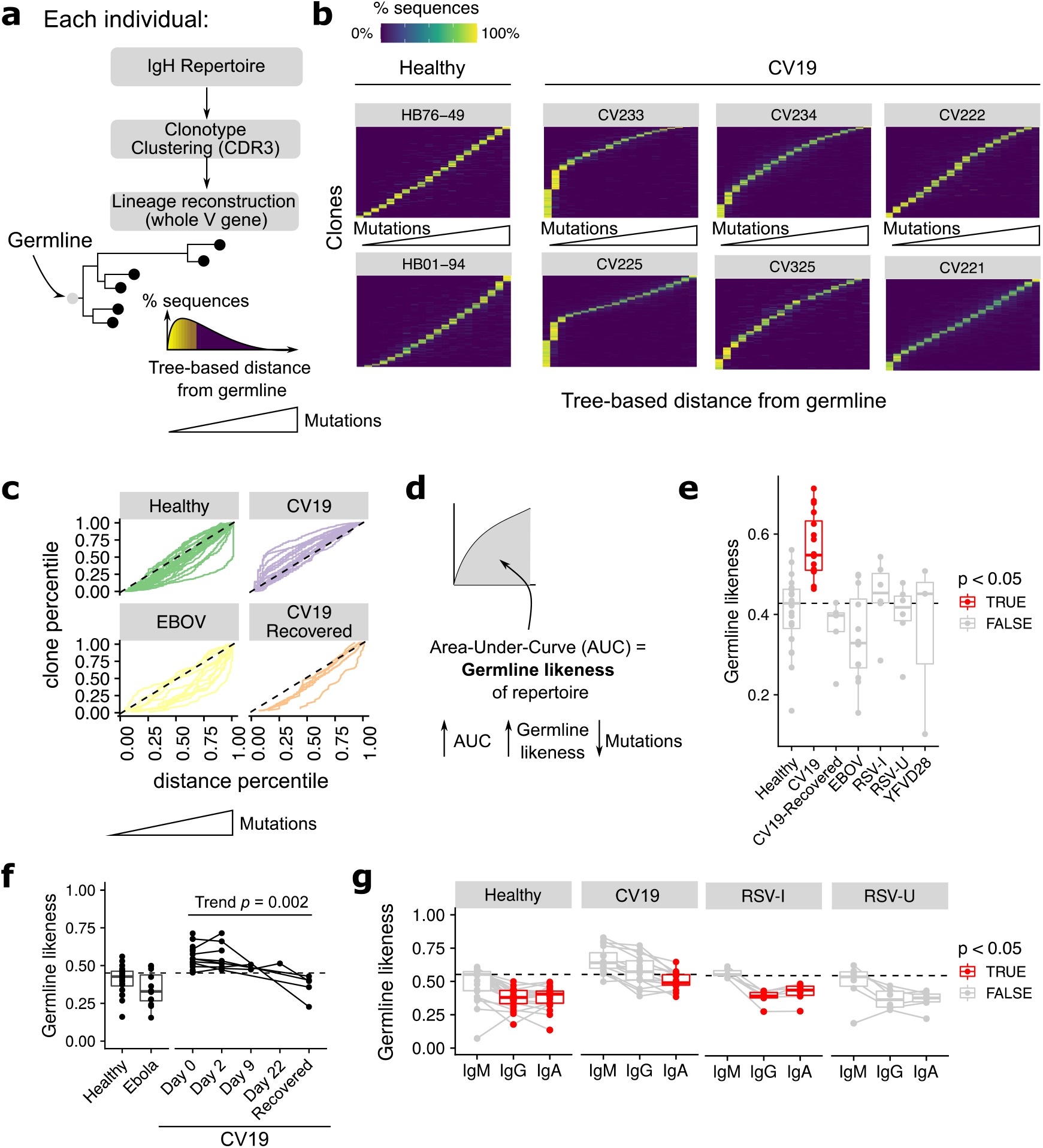
Mutational levels in BCR lineages. (**a**) Lineages trees were constructed by clonotyping the IgH CDR3 and the lineages reconstructed using the whole V gene rooting on the predicted germline, allowing the distance from germline to be estimated for each sequence. This allows ordering of sequences based on this distance from germline: depicted as a histogram [(**a**) bottom right]. (**b**) Clones in the repertoire, for selected donors, were ordered (vertical axis) using median distance from germline (horizontal axis), and the distribution of such distance for each clone was plotted with heatmap colours being the percentage of sequences within the clone containing the a given level of mutation. (**c**) Distance from germline distributions for every donor, split by condition, represented as curves. Dotted line represents the theoretical expectation of mutational level. (**d**) From each of these curves (in **c**) the area under the curve (AUC) was calculated giving a statistic of ‘Germline Likeness’, a higher AUC resembling more the germline and a lower AUC indicating more mutations. (**e**) Comparison of Germline Likeness between conditions: sample types with significant (p < 0.05) differences compared to Healthy (Wilcoxon rank-sum test) are highlighted in red with the dotted line being the healthy median. (**f**) The Germline Likeness across timepoints for CV19 patients with Healthy and Ebola data are reproduced here for comparison: trend was evaluated using the Jonckheere-Terpstra test. (**G**) Comparison of germline distance split by immunoglobulin isotype was performed split by cohort: significant (p < 0.05) differences compared to IgM (Wilcoxon rank-sum test) are highlighted in red.

### Ongoing Class switch recombination (CSR) detectable in PBMCs of CV19 and EBOV patients

Our lineage trees were further analysed for CSR events: respecting the sequential order of CSR in the genome, we identify CSR events where sequences of different antibody classes/subclasses are directly connected in the lineage tree. This enables us to trace the timing of CSR events (distance from the germline), the direction of class switching (e.g. from IgM to IgG1) and frequency of observation. Many clones have evidence of CSR in CV19 and EBOV, even after correcting for clone sizes (Figure 6a). In particular, CV19 patients were more likely to switch early to IgG1 from IgM, with little mutation (Figure 6b, c, d) and to IgA1 from either IgM or IgG1 later in the lineage with more mutation (Figure 6b,d). This agrees with the lack of CDRH3 “maturity” in IgG1 (Figure 2f) and the overall increased use of IgG1 and IgA1 seen in CV19 (Figure 2 a, b).

**Figure 6.**
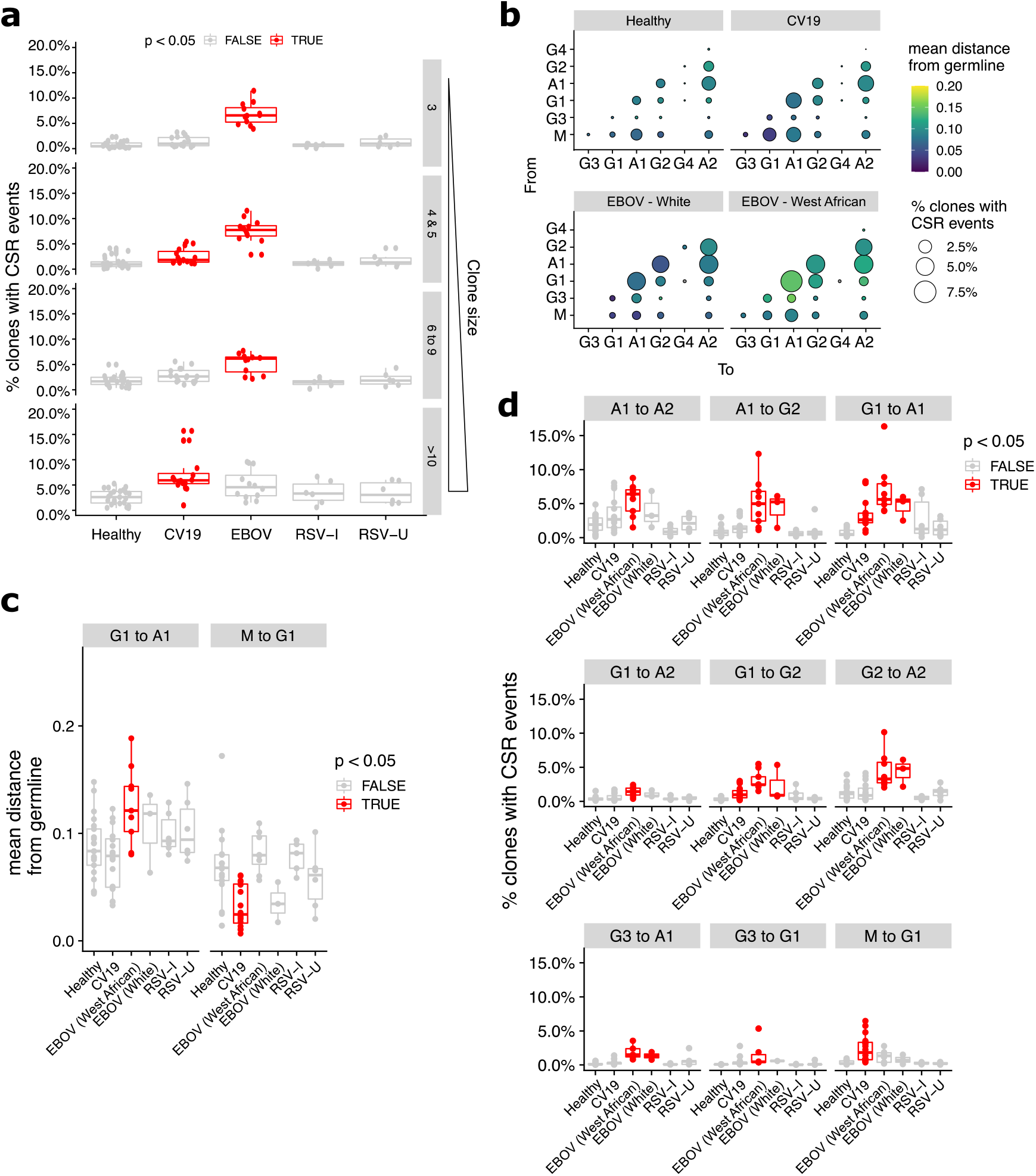
Prevalence of Class-switch recombination estimated from BCR lineage trees. (**a**) Lineage clones (see Figure 5) were assessed for prevalence of CSR events in terms of the proportion of clones and plotted by clone size and split by condition. (**b**) Bubble plot depicting the frequency and distance-from-germline of CSR events, separated by the CSR start (‘From’, vertical axis) and end (‘To’, horizontal axis) isotypes. Quantification was performed separately for different sample types. Bubble sizes are proportional to the frequency of CSR and colour is scaled by distance from germline at which CSR occurs, as estimated from the reconstructed lineage trees. (**c**) Statistical comparison of the median distance from germline at which CSR events occurred across sample types. Each donor was considered separately for every switch possibility. (**d**) Comparison of CSR frequency (percentage of clones with evidence of CSR) for each condition was also assessed for each donor (median, **d**). Statistical significance was evaluated using one-way ANOVA and Dunnett post-hoc comparison against Healthy with p < 0.05 highlighted in red (**c, d**). For (**d**), Supplementary Figure S8 contain analogous plots for all CSR combinations with significant (p < 0.05) differences compared against Healthy.

The evidence of increased CSR in convalescent EBOV patients is striking and occurs across the board with the exception of IgM switching to IgG1 (Figure 6d). We noticed that although there is a similar pattern of CSR preferences in White and West Africans individuals, the overall distance from germline is longer before CSR occurs in West Africans (Figure 6b). This may suggest that the ethnic bias in existing immunoglobulin sequence databases has resulted in mis-assignment of germline alleles. No CSR differences were seen in the RSV data.

## Discussion

We compared immunoglobulin gene sequences from pandemic (SARS-CoV-2), epidemic (Ebola) and endemic (Respiratory Syncytial Virus) patients in order to discover features that might distinguish newly emergent and endemic infections. The ability of B cells to generate a highly diverse immunoglobulin repertoire that might bind any antigen, and the diverse functionality of the antibodies produced, is critical for an effective immune response. Repertoire studies aim to identify specific antibodies by looking for biased usage of particular Ig genes, and have been useful in the past^16,27^. However, not all expanded genes encode specific binders^45^ and we need to consider the possibility that expansions found in the midst of an acute response may be a side effect of the disease involving inappropriate expansion of B cells carrying antibodies with off-target effects rather than a specific targeting to the challenge. Repertoire selection is normally a delicate balance between tolerance versus immune response to a pathogen and the inflammatory state of acute disease can upset the balance. Serological studies have shown an increase in autoreactive antibodies, particularly to interferons, during acute CV19 for example^50,51^.

Looking across different viral diseases, we found a general increase of *IGHV4-39* use in the repertoire of two different viral diseases (COVID19 and Ebola). Despite this, only one of our convergent clusters, dominated by CV19 sequences, use *IGHV4-39* (Figure 3c); it is possible that there are unannotated *IGHV4-39* SAR-CoV-2 binders. One single cluster does not, however, explain the larger expansion in *IGHV4-39* use across the COVID19 or Ebola repertoire. *IGHV4-39* may therefore be involved in the pathogenesis of the disease by promiscuous binding to self-proteins. Alternatively, *IGHV4-39* may simply support a wide range of specific binding properties, supported by the lack of convergence and given it has also been dominant in cancer, bacterial infection, influenza and HIV responses ^52–55^. Such promiscuous binders would have networks contributed to by more than 1 cohort with 52 networks matching this description in our data. It is also significant across all 64 large clusters 14 were dominated by CV19/CV19-R sequences yet only 5 matched known binders suggesting previously unknown SARS-CoV-2 specific binders.

In addition to expansion of gene use as an indicator of activation, we can infer biological information from assessment of the AID-mediated activities of CSR and SHM. These have long been associated with germinal centre formation ^56,57^. However, there is mounting evidence that CSR can occur prior to the germinal centre formation ^8–11^. T-independent activation has been shown to be driven by CD40-independant TLR/TACI activation ^58^. Our data indicates an early switching to IgG1 without extensive hypermutation. This data is consistent with Woodruff *et al* ^59^ who also found high germline similarity in IgA1, and ^39^ where CV19 samples were found to have more naïve-like characteristics. Our CV19 IgA1 sequences also indicate a lower level of hypermutation than the control group, albeit higher than the IgG1, likely reflecting their distance along the CSR hierarchy. Uniquely, our diversity analyses also indicate expansion of unmutated IgM clones. Alongside these data we see that CDRH3 region maturation of IgG1 and IgG3 genes in the CV19 patients is less removed from the IgM state than healthy IgG1 and IgG3, or any other class switched repertoires. Together with the lineage analysis of CSR timing, the whole picture in CV19 is of an early immature response of IgM, switching to IgG1 and then IgA1 but without much SHM, such as might occur in the absence of T cell help in a GC reaction. Whether these responses are unique to a live infection or because the virus is so novel is difficult to ascertain, with future vaccine and comparative studies likely to shed further light on this phenomenon.IgG1 is known for its antiviral properties ^46,60^ so is expected in this data. The majority of rapid immunological protection assays for COVID-19 focus on IgM or IgG ^61–64^. Since switching to IgA1 is notable in our data it would be useful for future serology assays to include IgA. Euroimmun’s IgA on LFA had one of the highest sensitivities at 87.8% compared to IgM and IgG from other assays (range 43.8-93%, mean 72.5%, median 76%) ^63,65^.

It is known that healthy older people generally have more antibodies capable of binding self-proteins^66^. The balance between antibodies with positive versus negative/bystander activity may be changed in older patients. We cannot tell this from our data except that we see a higher frequency of known spike binders clustering with CV19 repertoires in the younger cohort. Significant age-related differences occur in the dominant IGH CV19 genes: The increased use of *IGHV3-30* is only seen in older CV19 patients and that of *IGHV4-39* only in the younger group. We also see selection for *IGHD2-2, IGHD3-3* and *IGHJ6* only in the young, with *IGHJ6* occurring frequently in known binder networks, given the importance of *IGHD* and *IGHJ* genes to the CDRH3 region it is striking that the differences seen here are solely in the younger age group.

In comparison to our CV19 data our Ebola data paints an unusual picture where, even 2-3 months post-recovery with viral negative PCR tests, there are abnormally high proportions of class switched clones with little or no direction towards a particular sub-class. Given CSR is largely understudied, as far as we can tell such high rates of class switching, particularly so long after recovery, is entirely unique to this infection. Another unusual observation was that EBOV survivor’s memory B cell populations were more diverse than healthy controls suggesting stimulation with more diverse antigens, or a less structured and directed immune response. A ‘decay-stimulation-decay’ pattern resulting in the peak of antibody response being some 200 days after infection has previously been reported ^67^ and cytokine storms during infection may also be contributing to this phenomenon ^68,69^. It was not possible to collect blood samples from unrecovered patients, so we do not know if these observations were a requirement of patient recovery or a phenomenon unique to Ebola infection in general.

By comparing examples of pandemic, epidemic and endemic viral disease responses our results show that while aspects of B cell responses are unique to particular infections, the human immunoglobulin gene repertoire can show similarities of response across two very different diseases. There are many questions to be answered about the balance of beneficial versus bystander responses in acute inflammatory diseases, where the initial class switched responses seem to be immature (CV19) and possibly unregulated (EBOV infection). Coupled with the finding of genes such as *IGHV4-39* appearing in two completely different diseases, these data add weight to the hypotheses that an emergency humoral immune response to primary challenge can bypass normal stringent regulation and thus allow the production of autoimmune antibodies.

## Data Availability

IMGT/High-VQuest-annotated immunoglobulin sequence data file is available at Zenodo (https://dx.doi.org/10.5281/zenodo.5146019).

## Code Availability

Functions implemented to generate and analyse lineage trees were included in a R package BrepPhylo, which is available at https://github.com/Fraternalilab/BrepPhylo. All other code snippets used in analysing data presented in this work are available as R markdown files available at https://github.com/Fraternalilab/BrepPhyloAnalysis.

## Acknowledgements

This work was funded by the MRC (MR/L01257X/1 and MC_PC_15068), BBSRC (BB/T002212/1 and BB/V011456/1), EPSRC (EP/R031118/1) and UK National Core Studies (NCSi4P programme) funding. Ebola samples were kindly donated by Prof Alain Townsend (Oxford). Thanks also to the Surrey COVID-19 Collaboration and all volunteers for sample collection and to Gill Wallace and Vasiliki Tsioligka for technical support. The study “Convalescent plasma for early Ebola virus disease in Sierra Leone” (ISRCTN13990511 and PACTR201602001355272) was supported by the Wellcome Trust (Award 106491) and Bill and Melinda Gates Foundation; Public Health England Ebola Emergency Response; and the Blood Safety Programme, National Health Service Blood and Transplant. MGS was supported by the UK National Institute for Health Research Health Protection Research Unit in Emerging and Zoonotic Infections. We acknowledge the contribution of the Sierra Leone Association of Ebola Survivors and Ebola CP Consortium Investigators. The funders had no role in the collection and analysis of the samples, in the interpretation of data, in writing the report, nor in the decision to submit the paper for publication.

## Contributions

DDW, AS, JSO, MB designed study and protocols to collect and analyse COVID19 and YFV samples under REC 14/LO/1221

AS, MB, NK, ES, KL, CF, CCo, HL acquired and biobanked COVID19 samples CCh and PO provided RSV samples

MGS, JTS, JKB, AT provided Ebola samples

ES, AS, IS, JSO, AP, BW devised repertoire protocol and performed repertoire data generation

JN, AS, ES, DK, CM, DDW, FF performed bioinformatic analysis and data interpretation

All authors have read and commented on the manuscript

