## Supplemental Figures for "Pandemic, epidemic, endemic: B cell repertoire analysis reveals unique anti-viral responses to SARS-CoV-2, Ebola and Respiratory Syncytial Virus": Supplementary_Figures_20072021.pdf

### SUPPLEMENTARY MATERIALS

#### Supplementary Methods

| Primer | Application | Sequence (5'-3') |
|---|---|---|
| SmartNNN | TSO | AAGCAGUGGTAUCAACGCAGAGUNNNNUNNNNUNNNNUCTT[rG][rG][rG][rG][rG] |
| Smart20 | PCR1-Forward | CACTCTATCCGACAAGCAGTGGTATCAACGCAG |
| 10XSmart20 | PCR1-Forward | CACTCTATCCGACAAGCAGTCTACACGACGCTCTTCCGATCT |
| IGL-R1 | PCR1-Reverse | ACGGTGCTCCCTTCATGCGTG |
| IGK- R1 | PCR1-Reverse | GTAGTCTGCTTTGCTCAGCGTCAG |
| IGM- R1 | PCR1-Reverse | TGACGTCCTTGGAAGGCAGCAG |
| IGG- R1 | PCR1-Reverse | TCCTGAGGACTGTAGGACAGC |
| IGA- R1 | PCR1-Reverse | CTTCACGTGGCATGTCACGGAC |
| PID-Step | PCR2-Forward | [PID]CACTCTATCCGACAAGCAGT |
| IGL-R2 | PCR2-Reverse | [PID]TTCATGCGTGACCCGGCAGC |
| IGK- R2 | PCR2-Reverse | [PID]TGAGGCTGTAGGTGCTGTCCTTG |
| IGM- R2 | PCR2-Reverse | [PID]CGGGTACTGCTGATGTCAGAG |
| IGG- R2 | PCR2-Reverse | [PID]GACAGCYGGGAAGGTGTGCC |
| IGA- R2 | PCR2-Reverse | [PID]TACAGGTCCCCGGAGGCATC |

Supp. Methods Table 1: Primers used in amplification of B cell repertoire for template switch reverse transcription, gene specific PCR and semi-nested gene specific PCR with Patient Identifier (PID) multiplexing (See Supp. Methods Table 2 for PID sequences).

| Primer | Forward Sequence (5'-3') | Reverse Sequence (5'-3') | Library Usage |
|---|---|---|---|
| RSII-PID1 | GGTAGTCATGAGTCGACACTA | CCATCGCGATCTATGCACACG | EBOV, YFV, Healthy |
| RSII-PID2 | GGTAGTATCTATCGTATACGC | CCATCTGCAGTCGAGATACAT | EBOV, YFV, Healthy |
| RSII-PID4 | GGTAGACGTACGCTCGTCATA | CCATCTACAGCGACGTCATCG | EBOV, YFV, Healthy |
| RSII-PID9 | GGTAGTCATGCACGTCTCGCT | CCATCAGTATCACAGTCGCTG | EBOV, YFV, Healthy |
| RSII-PID17 | GGTAGCACGTCACTAGAGCGA | CCATCCAGACGTGACTGATAT | EBOV, YFV, Healthy |
| RSII-PID22 | GGTAGGTGCTGAGCATCAGAC | CCATCTGAGACATACTGAGTG | EBOV, YFV, Healthy |
| RSII-PID23 | GGTAGCACTGATCGATATGCA | CCATCATGTGCACTAGTGTAC | EBOV, YFV, Healthy |
| RSII-PID33 | GGTAGATACAGCACAGATGTG | CCATCGAGTCGTATCGCTCAT | EBOV, YFV, Healthy |
| RSII-PID44 | GGTAGCTCGATACGTGTAGCT | CCATCTGTCAGTAGATGACTC | EBOV, YFV, Healthy |
| RSII-PID45 | GGTAGGTGTCTAGACAGCTGT | CCATCTCGTACGAGATCGACA | EBOV, YFV, Healthy |
| Sequel_PID1 | GGTAGCACATATCAGAGTGCG | CCATCCACATATCAGAGTGCG | CV19, RSV, Healthy |
| Sequel_PID6 | GGTAGCATATATATCAGCTGT | CCATCCATATATATCAGCTGT | CV19, RSV, Healthy |
| Sequel_PID7 | GGTAGTCTGTATCTCTATGTG | CCATCTCTGTATCTCTATGTG | CV19, RSV, Healthy |
| Sequel_PID8 | GGTAGACAGTCGAGCGCTGCG | CCATCACAGTCGAGCGCTGCG | CV19, RSV, Healthy |
| Sequel_PID10 | GGTAGACGCGCTATCTCAGAG | CCATCACGCGCTATCTCAGAG | CV19, RSV, Healthy |
| Sequel_PID12 | GGTAGACACTAGATCGCGTGT | CCATCACACTAGATCGCGTGT | CV19, RSV, Healthy |
| Sequel_PID13 | GGTAGCTCTCGCATACGCGAG | CCATCCTCTCGCATACGCGAG | CV19, RSV, Healthy |
| Sequel_PID15 | GGTAGCGCATGACACGTGTGT | CCATCCGCATGACACGTGTGT | CV19, RSV, Healthy |
| Sequel_PID18 | GGTAGTCACGTGCTCACTGTG | CCATCTCACGTGCTCACTGTG | CV19, RSV, Healthy |
| Sequel_PID20 | GGTAGCACGACACGACGATGT | CCATCCACGACACGACGATGT | CV19, RSV, Healthy |
| Sequel_PID25 | GGTAGCGCGACACGCTCGCGC | CCATCCGCGACACGCTCGCGC | CV19, RSV, Healthy |
| Sequel_PID26 | GGTAGCACAGAGACACGCACA | CCATCCACAGAGACACGCACA | CV19, RSV, Healthy |
| Sequel_PID27 | GGTAGCTCACACTCTCTCACA | CCATCCTCACACTCTCTCACA | CV19, RSV, Healthy |
| Sequel_PID29 | GGTAGTATATATGTCTATAGA | CCATCTATATATGTCTATAGA | CV19, RSV, Healthy |
| Sequel_PID30 | GGTAGTCTCTCTATCGCGCTC | CCATCTCTCTCTATCGCGCTC | CV19, RSV, Healthy |

Supp. Method Table 2: Patient Identifier (PID) multiplexing sequences for the RSII and Sequel PacBio platforms and the libraries generated in which each was used.

**Supplementary Figures**

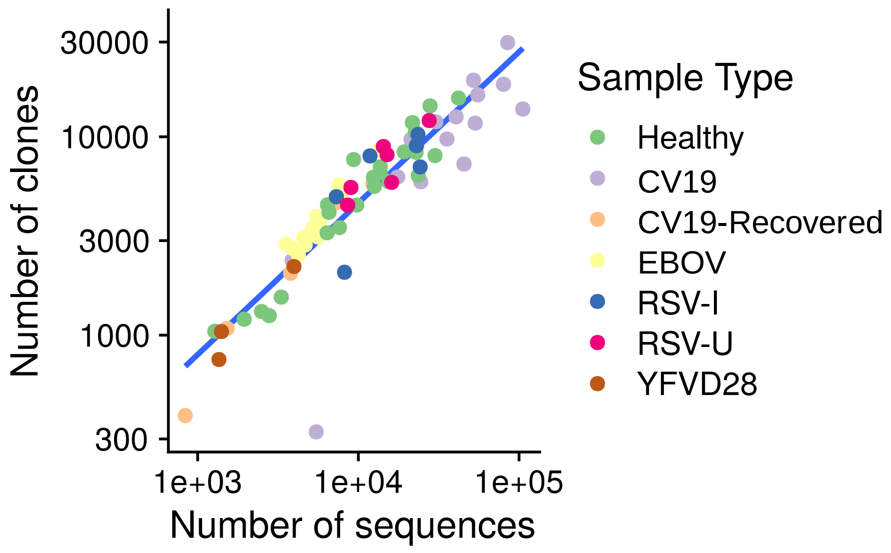


Figure S1. Relationship between the number of sequences sampled (horizontal axis) and the number of clones clustered (vertical axis) for each repertoire. Each data point represents repertoire from one donor, colour-coded by sample type.

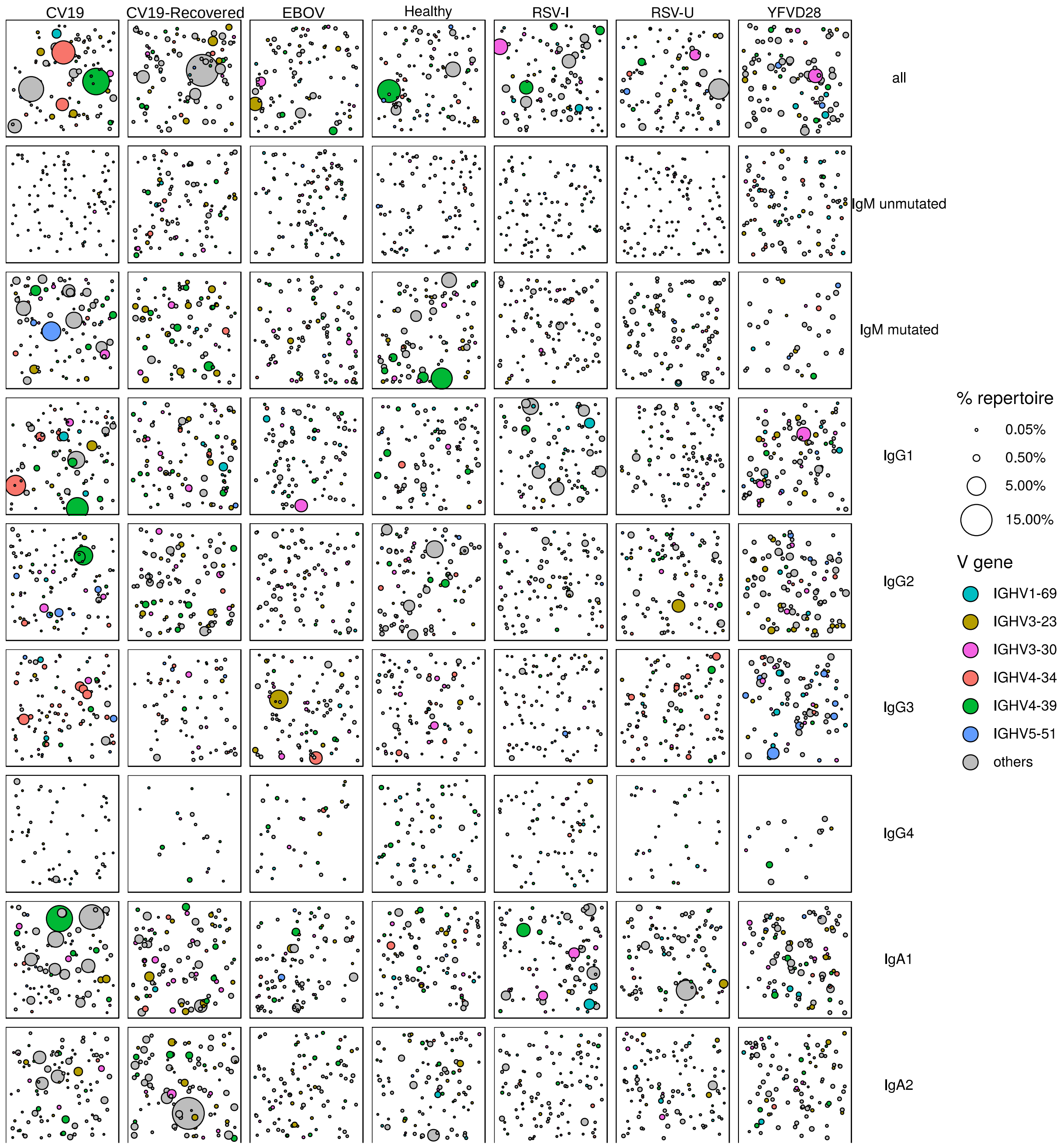


Figure S2. Clone size distribution in different cohorts and sequence subsets. Data identical to main text Figure 2C, except that here selected bubbles are colour-coded by V-gene usage discussed in the main text.

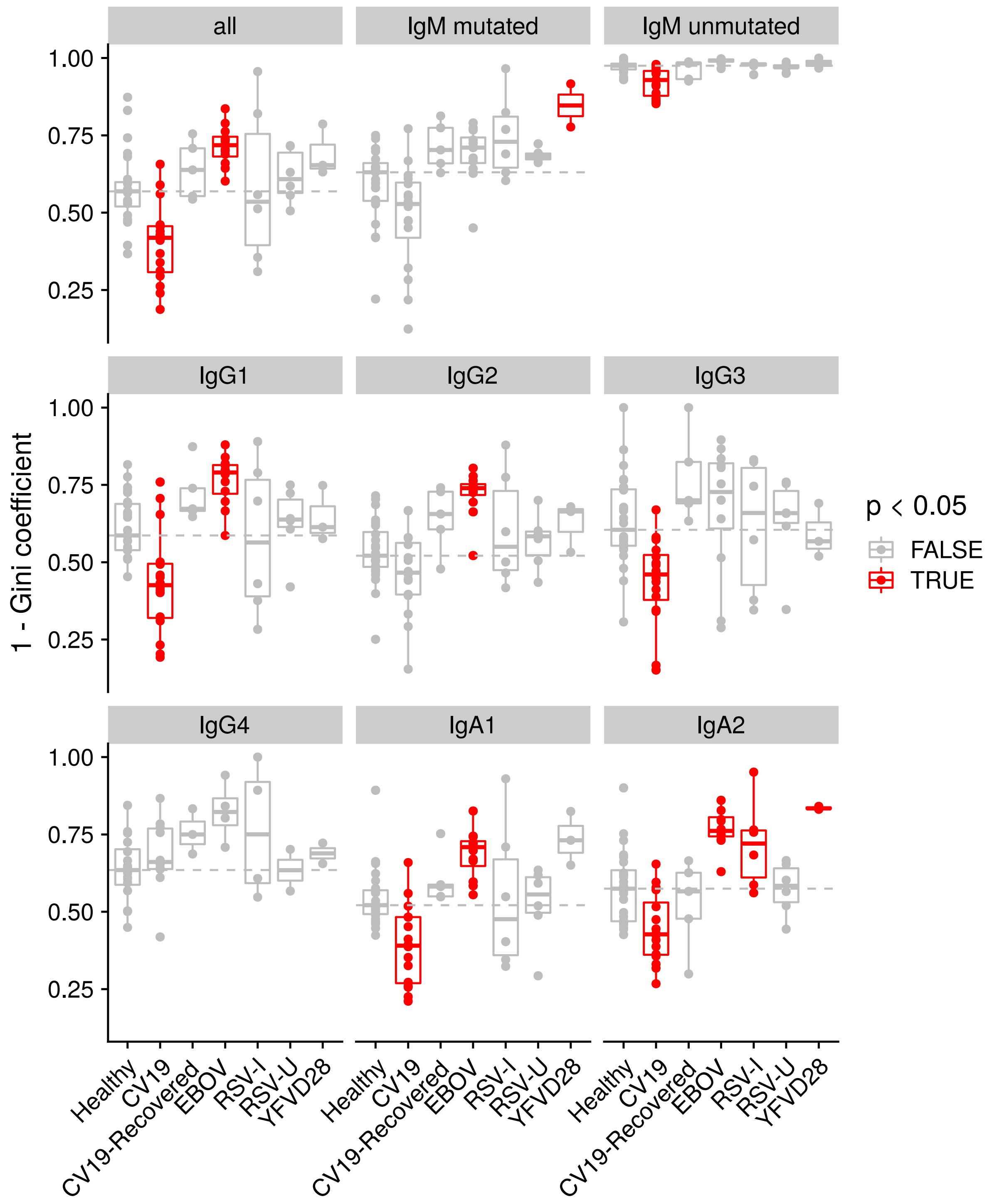


Figure S3. Diversity of clone distribution calculated using the formula (1 – Gini coefficient calculated on the clone size distribution). A larger value (closer to 1) indicates polyclonality and a smaller value (closer to 0) indicates monoclonality. Statistical significance was evaluated using a one-way ANOVA and Dunnett post-hoc comparison against the Healthy cohort; those with $p < 0.05$ were highlighted in red . Dashed line indicate the median diversity in the Healthy cohort as a reference. Selected panels are included as main text Figure 2D.

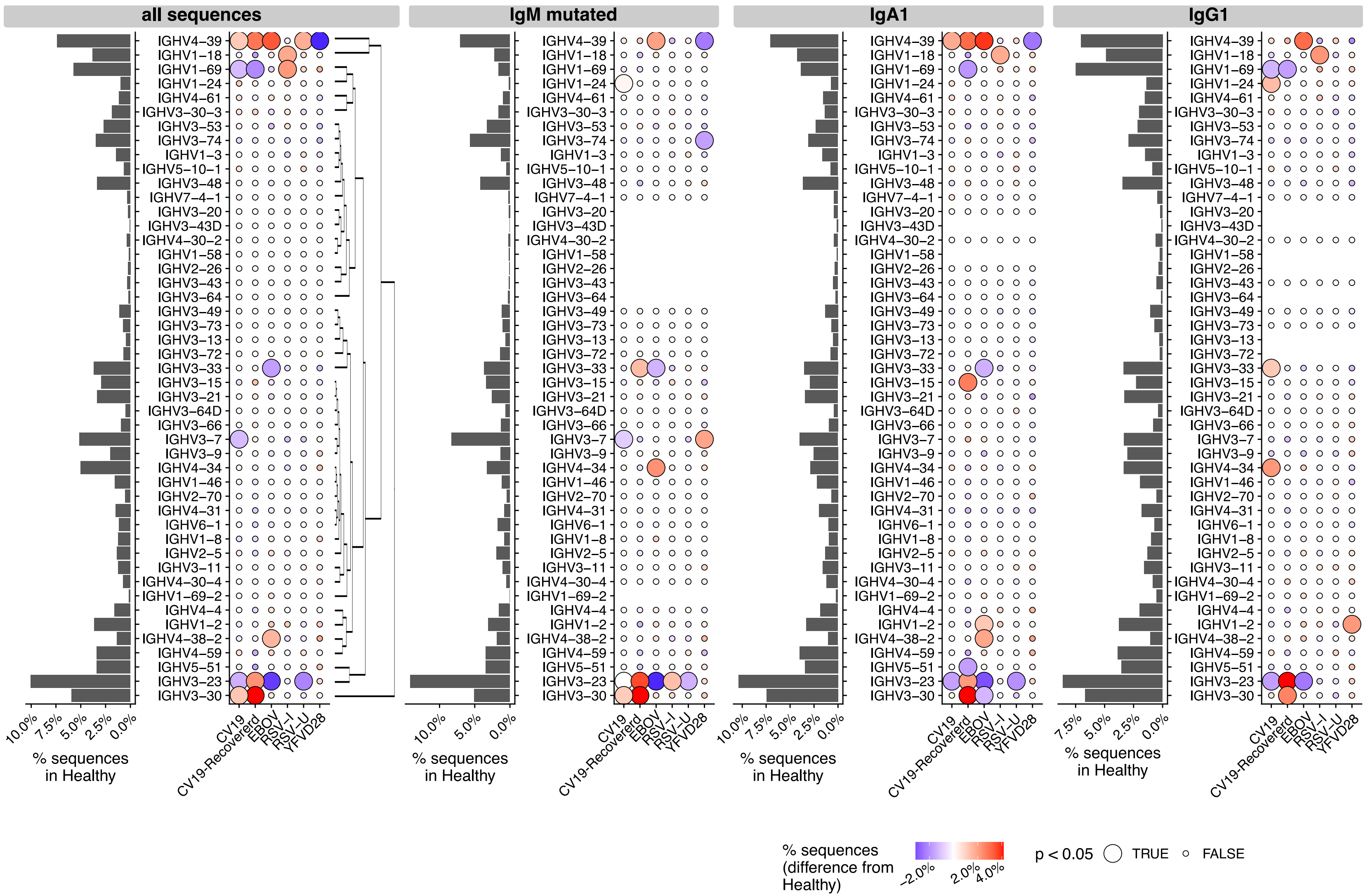


Figure S4. Usage of V genes (vertical axis) in Healthy (left, bar charts) and disease states (dot plots with colours and sizes representing difference and statistical significance in comparison to Healthy). Ordering of V genes is determined by hierarchical clustering of the V gene usage data from all sequences.

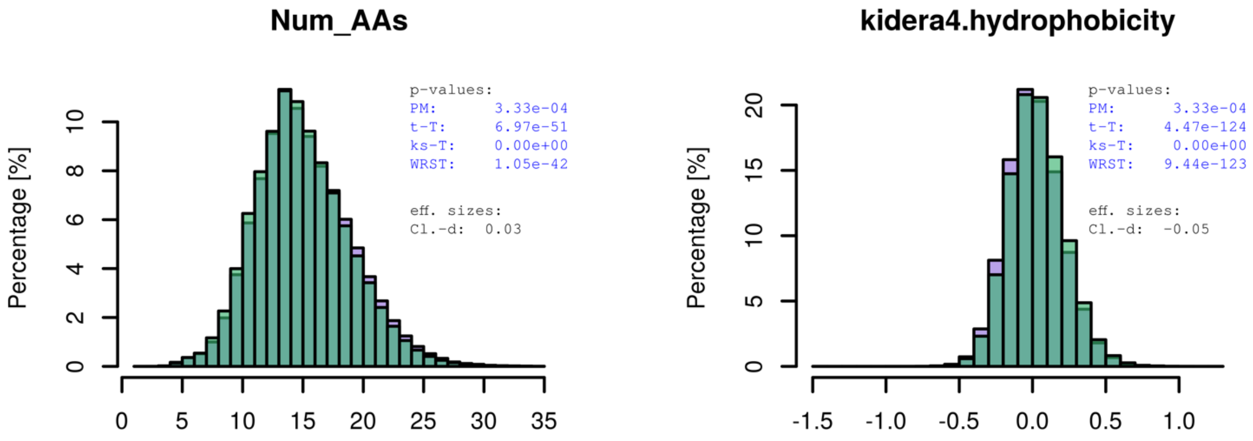


Figure S5. Histograms showing length (number of amino acids, left) and kidera factor 4 (hydrophobicity, right) distributions for CDR3 sequences from Healthy (green) and CV19 (lilac) repertoires. Here all sequences are considered without partitioning by isotypes and mutational level. Statistical comparisons and effect size (Cliff's delta) calculations were computed on the BRepertoire server (Margreitter et al, 2018).

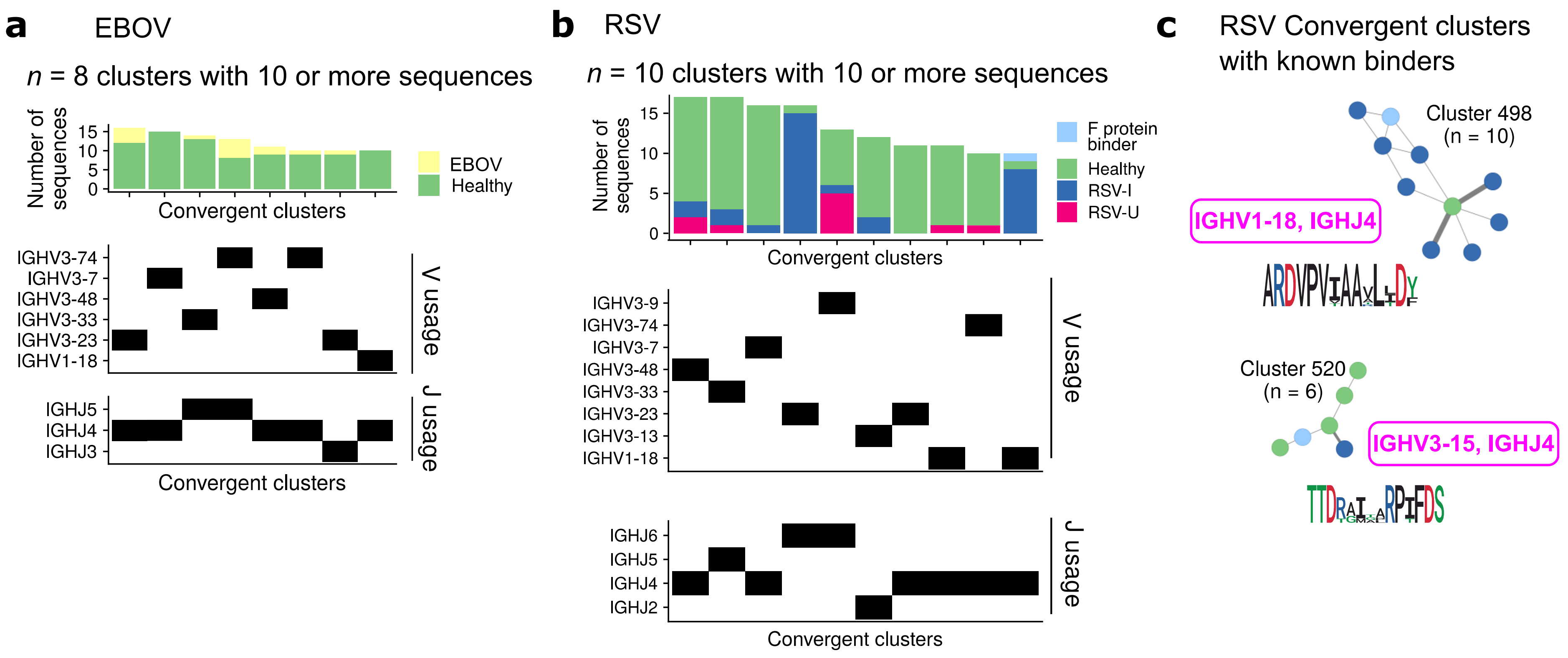


Figure S6. (**a,b**) Summary of convergent CDR3 sequence clusters for (**a**) EBOV and (**b**) RSV repertoire sequences. The construction of convergent CDR3 network is identical to what described in main text Figure 3A. Here shows the size and make-up (top, bar charts) of convergent sequence clusters with at least 10 sequences, as well as the V gene (middle, dark rectangle indicates this gene is used in the given convergent cluster) and J gene (bottom) usage. (**c**) Convergent clusters for RSV repertoire with known binders of the fusion glycoprotein ('F'). Colour scheme follows that of panel B. The V/J gene usage and CDRH3 amino acid sequences are shown.

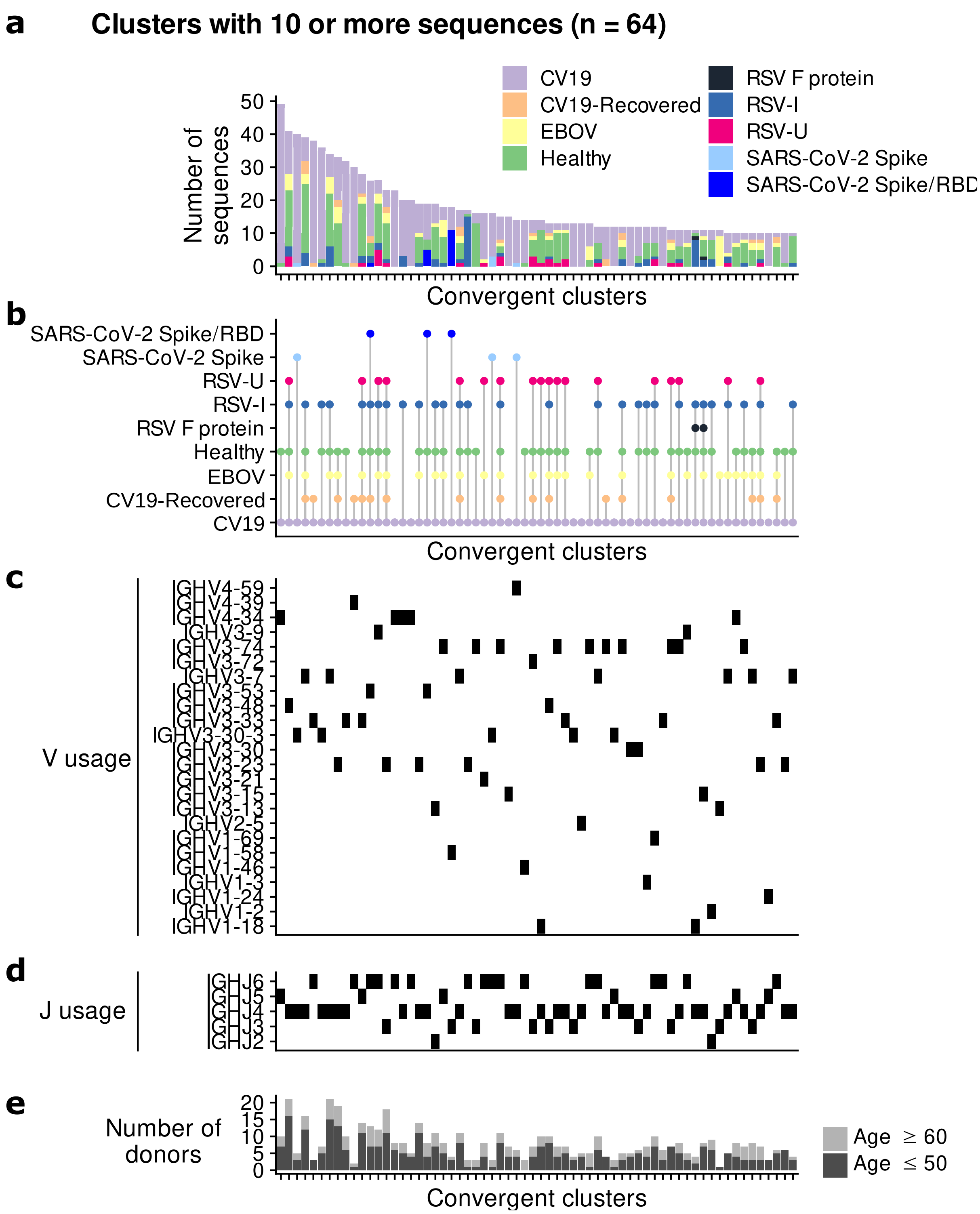


Figure S7. Convergent sequence clusters constructed considering CV19, EBOV and RSV repertoire and known binders altogether. This was performed to identify convergent clusters unique to a disease condition or universal across different conditions. Identical procedures as depicted in main text Figure 3A were followed. n = 64 clusters with more than 10 sequences were obtained. Panels **a, c, d, e** are identical to what shown in main text Figure 3C, depicting the breakdown (panel **a**), V gene usage (**c**) J gene usage (**d**) and number of donors represented (**e**) in each cluster. Panel B is an alternative representation of data in (**a**) with dots representing presence of different sequence types (vertical axis). This is provided to better highlight clusters shared across different sample types.

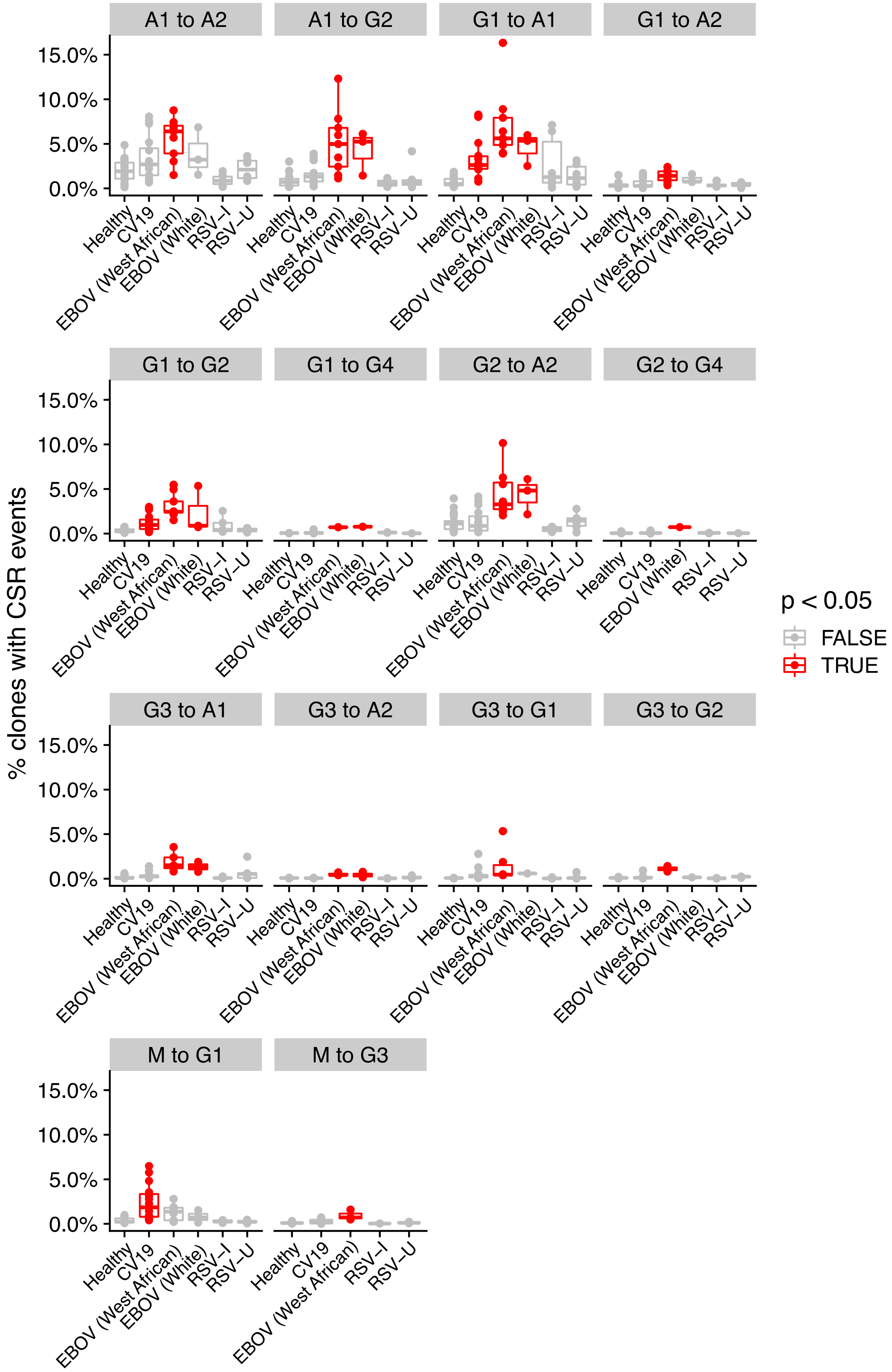


Figure S8. CSR Frequency (proportion of clones with CSR events) comparison across Healthy and disease states. Disease states with significant difference from Healthy (One-way ANOVA followed by Dunnett post-hoc comparison, $p < 0.05$) are highlighted in red. Data identical to that shown in main text Figure 6C except that here all CSR combinations with at least one comparisons with $p < 0.05$ are represented.

**Supplementary Tables**

| | |
|---|---|
| Table S1 | Donor characteristics. |
| Table S2 | Clone distribution. |
| Table S3 | Gene usage. |
| Table S4 | CDR3 characteristics. |
| Table S5 | Known antibody targeting SARS-CoV-2 proteins. |
| Table S6 | Sequences present in convergent binder networks. |
| Table S7 | Germline likeness of repertoires. |
| Table S8 | Frequency and distance-from-germline analysis of CSR events. |
